# A temporal record of the past with a spectrum of time constants in the monkey entorhinal cortex

**DOI:** 10.1101/688341

**Authors:** Ian M. Bright, Miriam L.R. Meister, Nathanael A. Cruzado, Zoran Tiganj, Elizabeth A. Buffalo, Marc W. Howard

## Abstract

Episodic memory is believed to be intimately related to our experience of the passage of time. Indeed, neurons in the hippocampus and other brain regions critical to episodic memory code for the passage of time at a range of time scales. The origin of this temporal signal, however, remains unclear. Here, we examined temporal responses in the entorhinal cortex of macaque monkeys as they viewed complex images. Many neurons in the entorhinal cortex were responsive to image onset, showing large deviations from baseline firing shortly after image onset but relaxing back to baseline at different rates. This range of relaxation rates allowed for the time since image onset to be decoded on the scale of seconds. Further, these neurons carried information about image content, suggesting that neurons in the entorhinal cortex carry information not only about when an event took place but also the identity of that event. Taken together, these findings suggest that the primate entorhinal cortex uses a spectrum of time constants to construct a temporal record of the past in support of episodic memory.

---

Episodic memory, the vivid recollection of an event situated in a specific time and place (Tulving, 1983), depends critically on medial temporal lobe (MTL) structures, including the hippocampus and entorhinal cortex (EC) (Milner, 1959; Eichenbaum, Yonelinas, & Ranganath, 2007; Dede, Frascino, Wixted, & Squire, 2016; Squire, Stark, & Clark, 2004). Building on pioneering work demonstrating a spatial code in the hippocampus and entorhinal cortex (O’Keefe & Dostrovsky, 1971; Fyhn, Molden, Witter, Moser, & Moser, 2004), recent research has shown that hippocampal representations also carry information about the time at which past events took place, suggesting that the MTL maintains a representation of spatiotemporal context in support of episodic memory (Pastalkova, Itskov, Amarasingham, & Buzsaki, 2008; MacDonald, Lepage, Eden, & Eichenbaum, 2011; Eichenbaum, 2017). Although a great deal is known about the temporal coding properties of neurons in the hippocampus, the temporal code in the entorhinal cortex, which provides the majority of the cortical input to the hippocampus is less understood, but see (Naya & Suzuki, 2011; Kraus et al., 2015; Naya, Chen, Yang, & Suzuki, 2017; Tsao et al., 2018).

Hippocampal time cells provide a record of recent events including explicit information about when an event occurred. Analogous to hippocampal place cells that fire when an animal is in a circumscribed region of physical space (Wilson & McNaughton, 1993; O’Keefe & Dostrovsky, 1971), hippocampal time cells fire during a circumscribed period of time within unfilled delays (Pastalkova et al., 2008; MacDonald et al., 2011; Kraus, Robinson, White, Eichenbaum, & Hasselmo, 2013). Across studies, there is a remarkable consistency in the properties of hippocampal time cells. Hippocampal time cells peak at a range of times during the delay interval and typically code time with decreasing accuracy as the delay unfolds, as manifest by fewer neurons with peak firing late in the delay and wider time fields later in the delay (Kraus et al., 2015; Salz et al., 2016; Mau et al., 2018). Hippocampal time cells have been observed in a wide range of behavioral paradigms, including tasks with and without explicit memory demands during the delay (Salz et al., 2016) and experiments in which the animal is fixed in space (MacDonald, Carrow, Place, & Eichenbaum, 2013; Terada, Sakurai, Nakahara, & Fujisawa, 2017). In addition, it has been shown that different stimuli trigger different time cell sequences (Terada et al., 2017; MacDonald et al., 2013). Taken together, time cells provide an explicit record of how far in the past an event took place, i.e., the amount of time that has passed since the beginning of a delay period or since the presentation of a to-be-remembered stimulus. By examining which time cells are active at a particular time, we can easily determine not only what event took place, but how far in the past that event occurred.

Many of the properties of hippocampal time cells have been observed in other brain regions including prefrontal cortex (Bolkan et al., 2017; Tiganj, Kim, Jung, & Howard, 2017; Tiganj, Cromer, Roy, Miller, & Howard, 2018; Jin, Fujii, & Graybiel, 2009) and striatum (Jin et al., 2009; Mello, Soares, & Paton, 2015; Akhlaghpour et al., 2016) suggesting that the hippocampus is part of a widespread network that carries episodic information. A recent report from the rat lateral EC adds important data to this growing body of literature regarding the representation of time in the brain. Tsao et al. observed a population of neurons that changed slowly and reliably enough to decode time within the experiment over a range of time scales (Tsao et al., 2018). Unlike time cells, which respond a characteristic time since the event that triggers their firing, lateral EC neurons respond immediately upon entry into a new environment, and then relax slowly. The relaxation times across individual neurons were very different, ranging from tens of seconds to thousands of seconds. To distinguish this population from time cells we will refer to neurons that are activated by an event and then relax their firing gradually as *temporal context cells*. The designation “temporal context cells” is not meant to indicate some intrinsic biological property but is simply intended, much like the terms “place cell” or “time cell” as a convenient short-hand to describe the functional properties of these neurons. Because these temporal context cells code for time, but with very different properties than time cells, these two populations provide a potentially important clue about the nature of temporal coding in the brain and the neural mechanisms that may support episodic memory.

Here, we identified temporal context cells in monkey EC during a free-viewing task (Meister & Buffalo, 2018). We examined EC neuron responses in a five-second period after presentation of an image. In the time after presentation of the image, a representation of what happened when should carry both time-varying information about when the image was presented, as well as information that discriminates the identity of the image. To anticipate the results, the data demonstrate that neurons in monkey EC are activated shortly after a visual stimulus and then decay with a variety of rates, enabling reconstruction of when the image was presented. This form of temporal coding is similar to temporal context cells observed in rat lateral EC (Tsao et al., 2018), but is qualitatively different from time cells that have been observed in the hippocampus and other regions. Because each image was shown twice over the course of the experiment, we were able to assess whether the pattern of activation over neurons depends on the identity of the image presented. Taken together, these data suggest that these temporal context cells carry information about what happened in addition to when it happened.

## Results

A total of 349 neurons were recorded from the entorhinal cortex (EC) in two macaque monkeys during performance of a visual free-viewing task. Each trial began with a required fixation on a small cross, followed by the presentation of a large, complex image that remained on the screen for five seconds of free viewing (Figure 1a). Unlike canonical hippocampal time cells, which are activated at a variety of points within a time interval (e.g., Figure 1c), most entorhinal neurons changed their firing a short time after the presentation of the image. Figure 2a shows three representative neurons that responded to image presentation (more examples are shown in Supplementary Figure S3). While most of these neurons increased their firing rate after the image was presented, some neurons decreased their firing rate in response to image presentation. Although behavior was not controlled during the five second free-viewing period, the response of these neurons was consistent across trials, which can be seen by examination of the trial rasters, indicating that the temporal responsiveness was unlikely to be a correlate of behavior.

**Figure 1.**
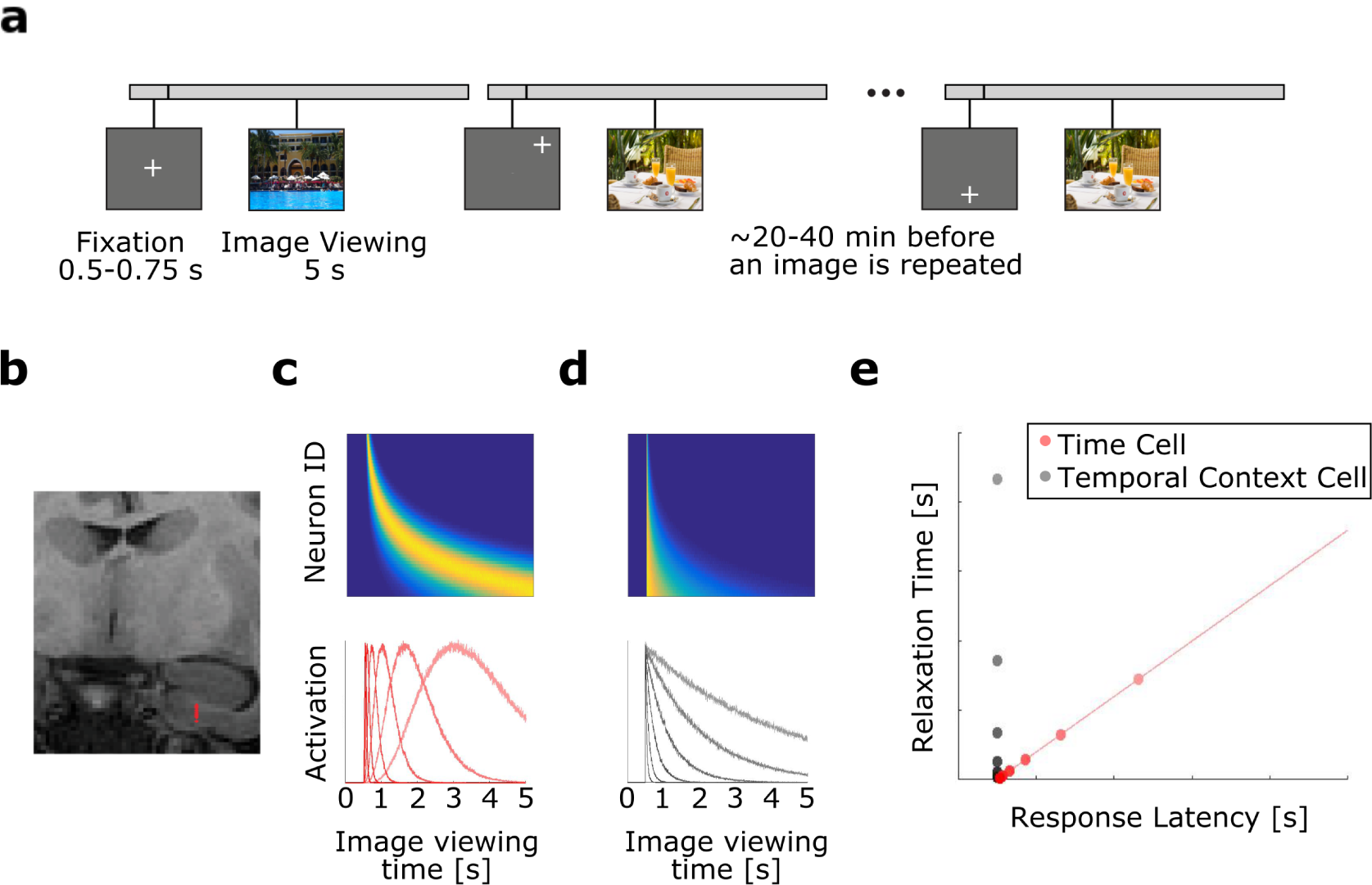
a-b: Summary of experimental procedures. **a**, Trial schematic for three trials. On each trial, the monkey freely viewed an image. After the monkey viewed an image for 5 seconds, the image disappeared. Following every trial (not shown), the monkey performed multiple gaze calibration trials and received a fruit slurry reward (see Methods for details). Images were presented twice during an experimental session. Between 20-40 minutes passed before an image was repeated. **b**, Estimated position of recording channels in the entorhinal cortex in one recording session is shown in red on a coronal MRI. **c-e: Two hypotheses for neural representations of time. c-d**, Heat plot (top) and tuning curves (bottom) for two hypotheses for how a time interval of image viewing could be coded in neural populations. In the heat plots, cooler colors correspond to low activity while warmer colors correspond to higher activity. **c,** Hypothetical activity for sequentially activated time cells, like those observed in the hippocampus. In this population, different neurons exhibit peak responses at different times indicating different firing fields. Because the time of peak response across neurons covaries with the spread of the firing field, neurons with later firing fields display wider firing fields. **d,** Hypothetical activity for monotonically decaying temporal context cells, like those observed in rodent EC. Neurons in this population reach their peak at about the same time. However, different neurons decay at different rates. **e**, Properties of neurons representing time passage by the hypothesis shown in **c** (red) or the hypothesis shown in **d** (gray). A population of time cells (red) should exhibit responses that occur at different times across a time interval, and these neurons should show a robust correlation between when response occurs and the time it takes the response to return to baseline. Conversely, a population of exponentially-decaying temporal context cells (gray) should exhibit responses that occur in a more restricted time range shortly after the start of a time interval, and these neurons should show no correlation between when peak response occurs and the time it takes to relax back to baseline.

**Figure 2.**
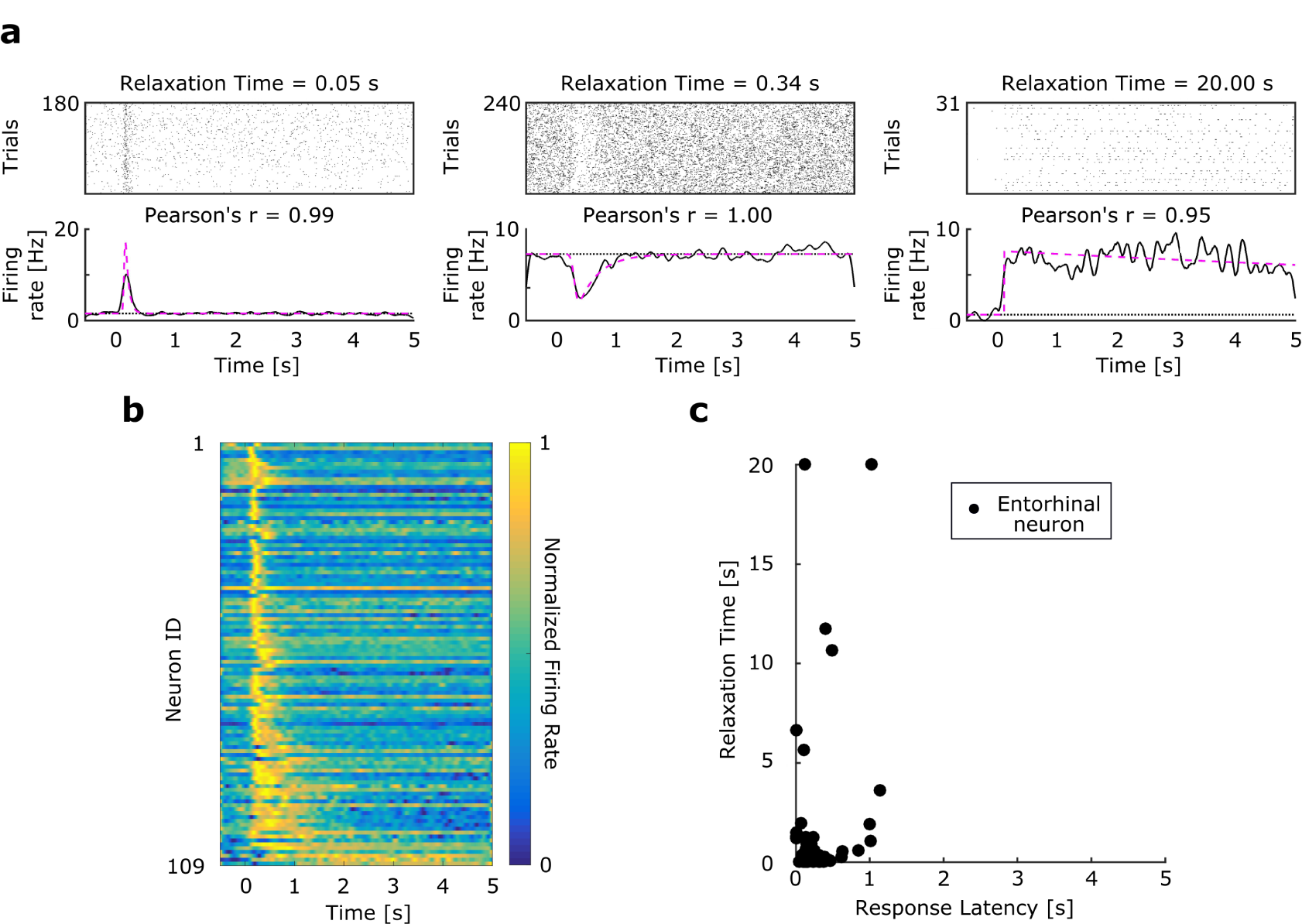
Temporal context cells in monkey entorhinal cortex respond shortly after image presentation, then return to baseline at a spectrum of rates. **a**, Three representative context cells that responded to image onset and decayed at different rates (Supplementary Figure S3 shows more examples). Each pair of plots indicate the activity of one neuron relative to image onset in a raster plot (top) and PSTH (bottom). In each raster, a tick mark indicates when the neuron fired an action potential. In the PSTH, the solid black line indicates the smoothed firing rate, and the pink line indicates the model estimate of firing rate. The estimated baseline firing rate is indicated by the black dotted line. Relaxation Time refers to the duration between response peak and when the neurons returned 63% of the way to baseline. Pearson’s *r* is the correlation between the fits for even and odd trials. **b**, A heatplot of the normalized firing rate of 109 temporal context cells, relative to image onset, sorted by their Relaxation Time. The color scheme is the same as in Figure 1c, d. The majority of neurons responded within 1 s of image onset, relaxing back to baseline with a spectrum of decay rates; some neurons relaxed back to baseline much sooner than other neurons that relaxed more slowly. **c**, A scatter plot of the joint distribution of each neuron’s Response Latency (the time at which a cell begins to respond) and Relaxation Time. Response Latencies did not span the entire 5 s, unlike Relaxation Time, and a neuron’s Response Latency and Relaxation Time were not correlated. Figure S2 shows the marginal distributions of these parameters.

Although the image-responsive neurons in EC responded at about the same time post-stimulus, they relaxed back to their baseline firing at different rates. Whereas some neurons relaxed back to baseline quickly (Figure 2a, left), some relaxed much more slowly. For instance the neuron shown in the right of Figure 2a did not return to baseline even after five seconds.

### Temporal receptive field analysis revealed a population of temporal context cells in entorhinal cortex

To quantify the qualitative results from examination of the individual units, we classified neurons that changed their firing in synchrony with image presentation and measured their temporal receptive fields with a model with three parameters. Each neuron’s firing rate in the time from 0.5 s before image onset to 5 s after was quantified using a model-based approach. The firing field was estimated as a convolution of a Gaussian (latency in neuronal response) and an exponential decay (Supplementary Figure S1a). This approach builds on previous work to estimate time cell activity as a Gaussian firing field (Tiganj, Kim, et al., 2017; Salz et al., 2016; Tiganj et al., 2018). The method for estimating parameters is described in detail in the methods. In this model, we were able to quantify apparent response properties using two key parameters: 1) the parameter *µ*, which describes the mean of the Gaussian, estimates the time at which each neuron begins to respond (Response Latency) and 2) the parameter *τ*, which describes the time constant of the exponential term, estimates how long each neuron takes to relax back to 63% of its maximum deviation from baseline firing (Relaxation Time) (Supplementary Figure S1b). A third parameter *σ* controls the standard deviation of the Gaussian term. Using this model, we tested whether a neuron had a time-locked response to image onset by quantifying the extent a model with a temporal response field fit the neuron’s data better than a model with only constant firing including the prestimulus period. Neurons that were better fit by including a temporal firing field were referred to as “visually responsive.” Importantly, this method would identify populations of either hippocampal time cells and temporal context cells as visually responsive. However, as shown in Figure 1e, these two different forms of temporal coding would produce distinguishable distributions of parameters.

### A substantial fraction of entorhinal neurons changed their firing in response to image presentation

In order to minimize the noise and obtain the most accurate distribution of response parameters across neurons, we used a conservative criterion to identify neurons that responded to image presentation. This method identified 109/349 neurons as visually responsive. Of those 109 responsive neurons, 84 neurons showed an increase in their firing rate in response to image onset, whereas 25 showed a decrease in their firing rate. Figure 2b summarizes the temporal response properties of these 109 neurons. Each row of the figure shows the averaged response of one neuron over the course of a trial. The data demonstrate that almost all of the neurons reached their maximum deviation from baseline within a few hundred milliseconds of the image presentation. This can be appreciated from the vertical yellow strip along the left edge of the heat map. These results are in striking contrast to the typical responses of hippocampal time cells (e.g., Figure 1c). Analogous plots for hippocampal time cells, which vary smoothly in their peak times, result in a curved ridge extending from the upper left to the lower right. In contrast, the variability across neurons in this entorhinal population was not in the time point at which the neurons reached their maximum deviation from baseline, but rather in the time course over which each neuron relaxed. This can be appreciated in the progressive widening of the ridge in Figure 2b from top to bottom.

### Visually responsive entorhinal units showed short Response Times but a broad distribution of Relaxation Times

Figure 2c shows the Response Latency and Relaxation Time for the 109 entorhinal neurons that were categorized as visually responsive (Supplementary Figure S2 shows the marginal distributions for each parameter). Response Latency values were clustered tightly at small values (median = 0.16 s, interquartile range = 0.13 s to 0.24 s). For 90% of neurons, the Response Latency was less than 0.40 s. In contrast, Relaxation Times showed a wider distribution (median = 0.23 s, interquartile range = 0.10 s to 0.61 s, 90^th^ percentile = 1.29 s), and even included values longer than the 5 s duration of the viewing period.

Lastly, the third parameter *σ*, which controls the standard deviation of the Gaussian, was small and tightly clustered across neurons (median = 0.02 s, interquartile range of 0.001 s to 0.06 s, 90^*th*^ percentile = 0.31 s). The consistently small value of this parameter indicates that the shape of the temporal receptive fields was well-described by a delayed exponential function.

### Response Time and Relaxation Time were not correlated across neurons

Across the 109 neurons, Response Latency and Relaxation Time were not significantly correlated with one another, Kendall’s *τ* = 0.03, *p* = 0.64. To assess whether this null effect was reliable, we computed the Bayes factor, which enables an estimate of the likelihood of the null hypothesis. This analysis yielded a Bayes factor of BF_01_ = 7.17, providing support that neuron Response Latency and Relaxation Time values are uncorrelated. Across neurons, Response Latency and *σ* were also not correlated with one another, Kendall’s *τ* = −0.04, *p* = 0.59, BF_01_ = 6.87. Unlike hippocampal time cells, there was no evidence that temporal context cells that peaked later in the time interval showed broader firing fields. In contrast to hippocampal time cells, which show a systematic relationship between the peak time of firing and the width of the temporal firing field (Kraus et al., 2013; Howard et al., 2014; Salz et al., 2016), the overarching conclusion from these analyses is that the firing of entorhinal neurons deviated from background firing shortly after the presentation of the stimulus and then relaxed exponentially at a variety of rates.

### Populations of entorhinal temporal context cells carry graded information about time

It is well-understood that hippocampal time cells can be used to decode the time since the beginning of a time interval (Mau et al., 2018) (Supplementary Figure S4). To assess the temporal information present in the population of entorhinal neurons, with special attention to the population of temporal context cells identified by the model-based analysis above, we trained a linear discriminant analysis (LDA) decoder to estimate time following presentation of an image. To the extent the predicted time bin for out-of-sample data is close to the actual time bin, one can conclude that the population response carried information about time. We first describe results from the entire population and then focus on the subpopulation of temporal context cells.

### Time was decoded better than chance from the population of entorhinal neurons

Figure 3 shows the results of the LDA on all 349 neurons from monkey EC. Our first question was whether or not the population contains information about time. For each time bin in Figure 3a, the confidence of the decoder (the posterior distribution) is shown across the range of possible time estimates. Perfect prediction would correspond to a bright diagonal; random decoding would correspond to a uniform gray square. Qualitatively, the non-uniformity of Figure 3a suggests that elapsed time can be decoded from the population of EC neurons. To quantitatively assess this, we found that the posterior distribution from the test data was reliably different from a uniform distribution using a chi-squared goodness of fit test, *χ*^2^(380) = 3906.8, *p* < 0.0001.

**Figure 3.**
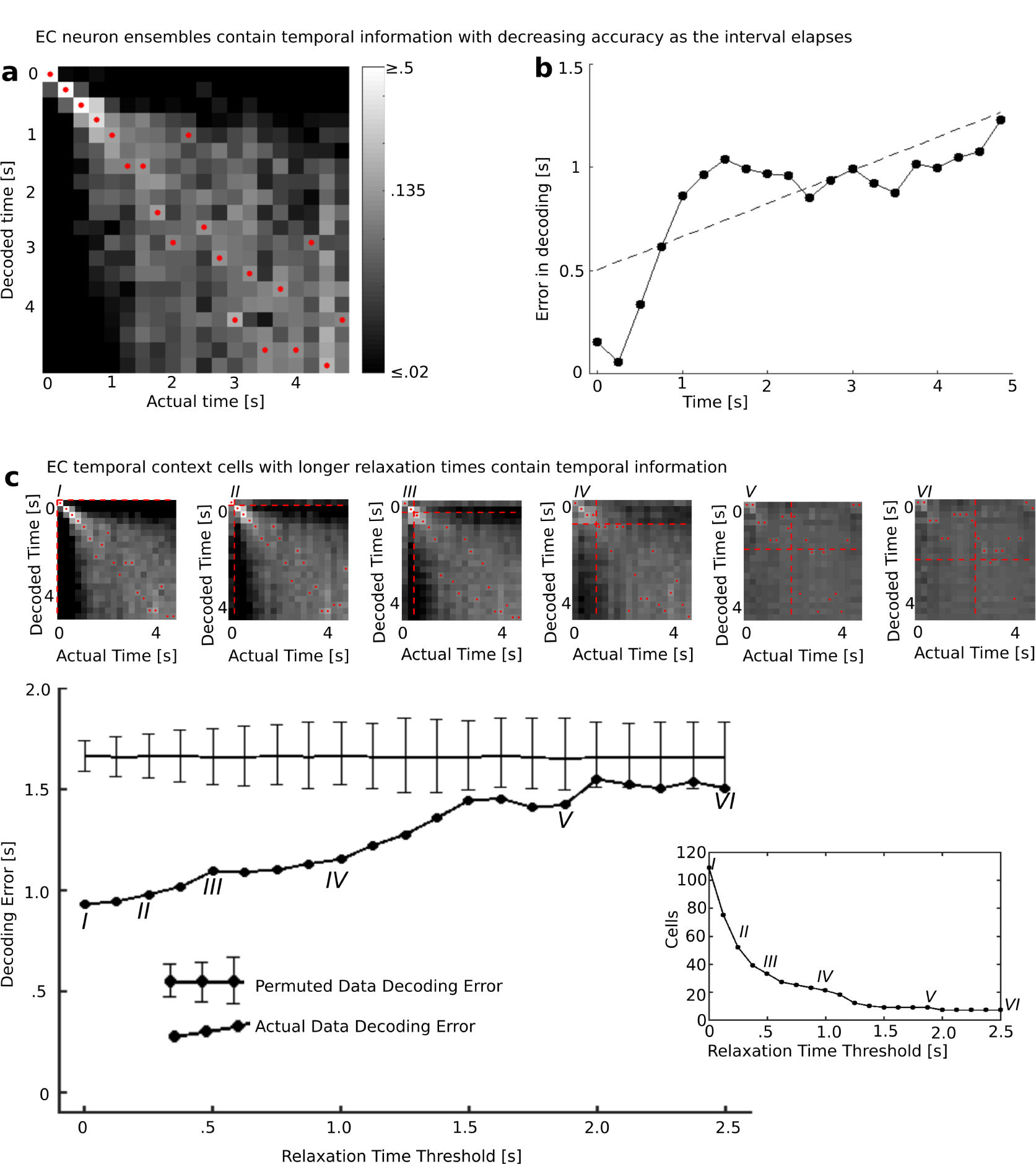
The population of entorhinal neurons encodes time with decreasing accuracy as the interval elapses. A linear discriminant analysis (LDA) decoder was trained to decode time since presentation of the image and then tested on excluded trials. **a,** Decoder performance trained on the entire population of EC neurons. The x-axis indicates the actual time bin which the LDA attempted to decode, the y-axis indicates the decoded time, and the shading indicates the log of the posterior probability, with lighter shading for higher probabilities (see color bar). The decoded time with the highest posterior for each actual time is marked with a red dot. The increasing spread of the diagonal and increasing dispersion of the red dots towards the lower right of the figure suggest that decoding accuracy decreases with the passage of time. **b,** Decoding error increases with the passage of time. The x-axis gives the actual time since image onset; the y-axis gives the mean of the absolute error produced by the decoder in **a**. The dotted black line is a fitted regression line. The absolute error increases with the passage of time. **c,** The temporal code was distributed along many temporal context cells, including those with slow relaxation times. The LDA decoder was first trained with the entire population of temporal context cells. To determine how the temporal code was distributed across the population of temporal context cells we re-ran the classifier but only allowing neurons with progressively longer Relaxation Times to contribute to the analysis. For a Relaxation Time Threshold of zero, all temporal context cells were included in the analysis, leading to results very comparable to the entire population (point labeled I). Then the Relaxation Time Threshold was increased from zero. For each value of the Threshold only temporal context cells with Relaxation Times greater than or equal to the Relaxation Time Threshold were included in the analysis. The large line graph shows absolute error averaged over all time bins as a function of Relaxation Time Threshold. Heatmaps showing the entire posterior distribution for selected points labeled by Roman numerals are shown along the top of the figure (same color bar and convention as in **a**). The threshold for relaxation time for each of the heatmaps is shown as dashed red lines. Note the earlier times which were most accurate (0-.75 seconds) dropped substantially in accuracy as the faster Relaxation Times were removed from the analysis. The horizontal line in the main linegraph shows the absolute error in decoding from a permuted dataset; error bars show the 0.025 and 0.975 quantiles. Right: the number of temporal context cells remaining in the analysis as a function of Relaxation Time Threshold. The gradual decline in accuracy suggests that temporal information was distributed smoothly throughout the population of temporal context cells.

Supporting this result, the mean absolute value of decoding error from the cross-validated LDA was reliably lower than the decoding error from training with a permuted data set. In each of 1,000 permutations we randomly reassigned the time bin labels of the training events used to train the classifier. The absolute value of the decoding error for the original data was 0.923 s, which was more accurate than the mean absolute value of the decoding error for all 1,000 permutations. As shown in Supplementary Figure S5a, the values for the permuted data were approximately normal with mean 1.65 s and standard deviation 0.04 s, resulting in a z-score of more than 18 (*z* = 18.175). These analyses demonstrate that time since image presentation could be decoded from populations of neurons in monkey EC.

### The precision of the time estimate decreased as the interval unfolded

Although the population response in entorhinal cortex could be used to reconstruct time, inspection of Figure 3a suggests that the precision of this reconstruction was not constant throughout the interval. Figure 3b shows the average absolute value of the decoding error at each time bin. These data suggest that this error increased as a function of time. A linear regression of decoding error as a function of time showed a reliable slope, 0.16 ± 0.03, as well as intercept 0.55 ± 0.1, both *p* < 0.001, *R*^2^ = 0.56, *df* = 18. The information in the entorhinal population about the time of image presentation decreases in accuracy as the image presentation recedes into the past.

### Time can be decoded well past the peak firing of temporal context cells

Theories that proposed the existence of temporal context cells argue that they convey information about time *via* their gradual decay. Another possibility is that the temporal context cells only carry decodable information about time because of their rapid deflection near time zero. If that is the case, then the population of entorhinal neurons should only carry information about time in the period of time close to the Response Time. To assess how far into the interval time could be reconstructed, we repeated the LDA analysis excluding progressively more time bins starting from zero. If the LDA can reconstruct time above chance using only bins corresponding to times ≥ *t*, then we can conservatively conclude that time can be reconstructed at least time *t* into the interval. To assess this quantitatively, the actual data were compared with permuted data for each repetition of the LDA using absolute error to assess performance (see Methods for details, Supplementary Figure S5). This analysis showed that time more than 2.25 s after the image onset can be reliably decoded (*p* < 0.01). This conservative estimate is an order of magnitude longer than the median value of the peak time (0.160 s), suggesting that the gradual decay of temporal context cells could be used to reconstruct information about time. Decoder performance varies later into the delay, with the performance of the decoder actually improving as noisier bins are eliminated from the analysis. For instance, time at 3 s can be reliably decoded (*p* < .01). The decoder is last significantly better than chance at *p* < 0.01 with 3.0 seconds excluded (below the line marked with ‘**’).

### Information about time was distributed throughout the population of temporal context cells

Theories that proposed the existence of temporal context cells argue that they convey information about time *via* their gradual decay. To determine how temporal information was distributed across the population of temporal context cells we performed a decoding analysis restricting our attention to temporal context cells. The analysis was performed initially using all temporal context cells, and then progressively removed cells with Relaxation Times shorter than a Relaxation Time Threshold. The Relaxation Time Threshold ranged from 0 s—including all 109 temporal context cells—to 2.5 s—at which point only 7 temporal context cells remained in the analysis. For each Relaxation Time Threshold, performance was summarized by averaging decoding error across all bins. If only a subpopulation of temporal context cells with fast Relaxation Times contributed to the temporal information in the ensemble, we would expect an abrupt decrease in performance as the Relaxation Time Threshold passed through that critical value.

These results are shown in Figure 3c, with selected decoder posteriors shown for various Relaxation Time Thresholds (Figure 3c*I-VI*). The population at each Relaxation Time Threshold was used to generate its own permuted data set. The first observation is that the performance of the decoder changed gradually, suggesting that temporal context cells with a range of Relaxation Times conveyed useful information about the time of image presentation. Examination of the heatmaps in Figure 3c suggests that excluding temporal cells with Relaxation Times below a particular value (indicated by dashed red line) disrupts the ability to distinguish times below that value. However, the ability to decode time above the threshold is relatively intact. Statistically, the decoder performed better than chance even with a Relaxation Time Threshold of 1.875 s (with 9 cells remaining). This analysis suggests that temporal information is distributed throughout the population of temporal context cells. Further the population conveys information about a range of time scales because the population has a variety of Relaxation Times.

### EC neurons conveyed information about image identity

In this experiment, each image was presented twice. Although it was not practical to assess image coding using a classifier, it was possible to exploit the repetition of images to determine whether EC neurons contained information about image identity. This question was addressed using both single cell analyses and population analyses, which showed convergent results. In both cases we compare the first and second presentation of the same image to the first and second presentations of different images. Note that because these analyses always compare a first presentation to a second presentation, they are not confounded by repetition effects that have been observed in entorhinal neurons (Xiang & Brown, 1998; Meyer & Rust, 2018; Jutras & Buffalo, 2010).

### Firing rate of individual neurons was correlated for same presentation of images

For each neuron we assembled an array giving the firing rate during the first presentation of each image (averaged over 5 s) and asked whether this array was correlated with the firing rate of second presentations of the same images. Many individual EC neurons responded to several images, as reported in more detail in the Supplemental text (see especially Supplementary Figure S7). If the firing rate of a neuron depends on the identity of the image, we would expect to see a positive correlation using this measure. For this analysis we restricted our attention to the neurons (*n* = 270) in the entorhinal population that were recorded long enough to be observed for both first and second presentations of a block of stimuli (repetitions were separated by 20-40 minutes).

The mean correlation coefficient (Kendall’s *τ*) across neurons was significantly greater than zero, *τ* = 0.06 ± 0.02, *t*(269) = 7.69, *p* < 0.001, Cohen’s *d* = 0.47, indicating that the spiking activity of many neurons depended on image identity (Supplementary Figure S6b). This comparison was confirmed by a Wilcoxon signed rank test on the values of Kendall’s *τ, V* = 27577, *p* < 0.001. This finding was also observed for the subset of visually responsive neurons (*n* = 93) that we describe as temporal context cells. Taken in isolation, the temporal context neurons showed a mean correlation coefficient significantly greater than zero, as measured by t-test, 0.09 ± 0.02, *t*(89) = 7.34, *p* < 0.001, Cohen’s *d* = 0.77 and Wilcoxon signed rank test, *V* = 3553, *p* < 0.001. Neurons that were not temporal context cells (*n* = 179) also had a mean correlation coefficient different from zero, 0.04 ± 0.02, *t*(176) = 4.65, *p* < 0.001, Cohen’s *d* = 0.35, *V* = 11363, *p* < 0.001. These results are consistent with a population that contains information about stimulus identity.

### The population of EC neurons was more similar for repeated presentations of the same image

The preceding analyses show that the firing of many EC neurons distinguished image identity above chance. If the response of the entire population contained information about stimulus identity, we would expect, all things equal, that pairs of population vectors corresponding to presentations of the same image would be more similar to one another than pairs of population vectors corresponding to presentations of different images. To control for any potential repetition effect, these analyses compared the similarity between the repetition of an image with its original presentation to the similarity between the second presentation of an image with the first presentation of a different image. To control for any potential recency effects, we swapped images adjacent to the original presentation of the target image. To be more concrete, if we label a sequence of images initially presented in sequence as A, B, and C, we would separately compare the population response to the repetition of B to the response to the initial presentation of A, B, and C. We refer to the similarity of the second presentation of B to the first presentation of B as lag 0. The similarity of the second presentation of B to the first presentation of A is referred to as lag −1; the similarity to the first presentation of C is lag +1. To the extent that the similarity at lag 0 is greater than lag −1 and +1, we can conclude that the population vector distinguishes image identity.

Supplementary Figure S6d shows the results of this population analysis. The similarity for lag 0 pairs (comparing population response to an image with its repetition) was greater than the similarity for lags ±1 (comparing the response to neighbors of its original presentation). Statistical comparisons to lags ±1 each showed a reliable difference. A paired t-test comparing population similarity at the level of blocks (*n* = 64 trial blocks of repeated images) showed that similarity at lag 0 was reliably larger than both lag +1, 0.012 ± 0.005, *t*(63) = 5.11, *p* < 0.001, Cohen’s *d* = 0.64, and lag −1, 0.014 ± 0.006, *t*(63) = 5.02, *p* < 0.001, Cohen’s *d* = 0.63. To evaluate the same hypothesis using a non-parametric method, we performed a permutation analysis by randomly swapping within-session pairs of lag 0 and lag ±1 and calculating the mean difference between the pairs 100,000 times. The observed value exceeded the value of 100,000/100,000 permuted values for both lags +1 and −1. We conclude that the population response was more similar for presentations of the same image than for presentations of different images. This analysis, which controlled for repetition and recency, demonstrates that the response of the population of EC neurons reflected image identity. Coupled with the other results reported here, this means that the population carried information about what happened when.

## Discussion

Episodic memory requires information about both the content of an event as well as its temporal context (Tulving, 1983; Eichenbaum et al., 2007; Eichenbaum, 2017). In this study, many EC neurons responded to the onset of the image. These *temporal context cells* responded to image onset at about the same time, within about 300 ms of image onset. However, different temporal context cells showed variable rates of relaxation back to baseline (Figure 2). Information about time since the image was presented could be decoded due to gradually decaying firing rates over a few seconds (Figure 2d-e). Notably, the relaxation rate was not constant across neurons, but rather showed a spectrum of time constants. The population vectors following repeated presentations of the same image were more similar to one another than to presentations of different images. This, coupled with several control analyses, show that the firing of entorhinal neurons also distinguished stimulus identity. Taken together, the results demonstrate that in the time after image presentation, the population of EC neurons contained information about what happened when.

Sequentially activated time cells, such as have been observed in the hippocampus (Pastalkova et al., 2008; MacDonald et al., 2011; Salz et al., 2016), medial entorhinal cortex (Kraus et al., 2015) and many other brain regions (Jin et al., 2009; Mello et al., 2015; Tiganj et al., 2018; Tiganj, Shankar, & Howard, 2017) also contain information about what happened when in the past. However, entorhinal temporal context cells have very different firing properties than sequentially activated time cells. As a population, time cells convey the amount of time that has passed since the occurrence of an event by firing at different temporal delays after the triggering event. In contrast, EC temporal context cells all responded at about the same time but relaxed at different rates. The range of relaxation times enables the population to convey information about different time scales. For instance, a temporal context cell that returns to baseline firing within 1 second would not be effective in distinguishing a 5 second interval from a 10 second interval. In contrast, a cell that decays back to baseline around 7 seconds would be effective in distinguishing this longer time period. In this way, a range of decay rates enables the population of temporal context cells to decode time over a wide range of time scales.

### Relationship to findings from rodent and monkey EC

The pattern of results observed here aligns well with a recent report from rodent lateral EC (Tsao et al., 2018). In that study, lateral EC neurons changed their firing in response to a salient event, i.e., the animal entering a new environment, and then relaxed back to baseline monotonically. Notably, different neurons relaxed at different rates with time constants ranging from tens of seconds to many minutes. However, despite the many methodological differences between that study and this one—rats moving through a series of open enclosures *vs* seated monkeys observing a series of images—the response properties shared striking similarities, suggesting a common computational function for EC across species.

The results in this paper are consistent with studies that have shown long-lasting responses in EC neurons in different preparations in rodents. For example, sustained responses have been observed in both in-vitro (Klink & Alonso, 1997; Egorov, Hamam, Fransén, Hasselmo, & Alonso, 2002; Tahvildari, Fransén, Alonso, & Hasselmo, 2007; Yoshida, Fransén, & Hasselmo, 2008; Hyde & Strowbridge, 2012) and anesthetized (Hahn, McFarland, Berberich, Sakmann, & Mehta, 2012; Leitner et al., 2016) approaches. Similarly, slow changes in firing rate were observed in the EC of rats during trace eyeblink conditioning across distinct environmental contexts (Pilkiw et al., 2017). Computational modeling studies have suggested that properties of a calcium non-specific cation current observed in slice are sufficient to generate a spectrum of response decay periods, ranging from brief to prolonged (Tiganj, Hasselmo, & Howard, 2015; Liu, Tiganj, Hasselmo, & Howard, 2019). Juxtaposed with work showing spatial responses in navigating rodents (Deshmukh & Knierim, 2011; Wang et al., 2018; Høydal, Skytøen, Andersson, Moser, & Moser, 2019), these findings suggest EC neurons code for “position” in both temporal and spatial domains.

The current results are also consistent with previous findings from the monkey medial temporal lobe. A recent investigation of single-neuron activity across shorter timescales (within a ∼1 s delay) identified time-varying responses in the hippocampus, but not the entorhinal cortex (Naya et al., 2017). However, in earlier work with longer trial durations, entorhinal neurons exhibited response dynamics across several seconds (Suzuki, Miller, & Desimone, 1997). The current results also mirror previous studies showing coding for for temporal information in monkey PFC attributable to slow ramping activity (Machens, Romo, & Brody, 2010; Rossi-Pool et al., 2016, 2019). It remains to be seen if those findings reflect similar or distinct computational mechanisms to those observed in EC in this study.

### Exponentially-decaying neurons with a spectrum of time constants is the Laplace transform of time

Why would the brain use two distinct coding schemes—time cells versus temporal context cells—to represent time? One proposed answer is that there might be a local circuit processing advantage from having time cells that can signal a specific moment at the single-neuron level, instead of having that information exist in the collective responses of different temporal context cells. Mechanistically, the brain may achieve creating a time cell response by combining the responses of temporal context cells (Shankar & Howard, 2012). The mathematical description of this process would begin with the brain estimating a temporal record of the past as neural activity that is the real Laplace transform of a function of past time (Shankar & Howard, 2012, 2013; Howard et al., 2014).

Cells coding for the real Laplace transform have exponential receptive fields with a variety of rate constants, very much like the results observed here in entorhinal neurons (Figure 2). In this proposal, the activity representing the contents of the past changes in the time after an image presentation. At a time *t* after image presentation, the neural representation of the past is a function with the presentation of the image at time *t*. As *t* increases, this function changes smoothly. A population of neurons coding the real Laplace transform of time should thus change shortly after image presentation and then relax exponentially, with different neurons relaxing at different rates.

Time cells like those in the hippocampus, in contrast, can be described by performing an additional computation upon the activity described above. The inverse Laplace transform — which can be approximated using a set of feed-forward connections with center-surround weights on the activity described above — directly estimates what happened when. Instead of exponential receptive fields, cells coding for the inverse transform have circumscribed receptive fields that tile the time axis. As the image presentation recedes into the past, neural response to its presentation resembling the inverse transform would generate a series of sequentially-activated time cells.

### Laplace transforms of other variables in the MTL

This computational framework for representing functions over continuous variables using the Laplace transform and its inverse can be generalized from time to other variables as well (Howard et al., 2014). For instance, border cells in EC (Boccara et al., 2010) code for distance from an environmental landmark with monotonically-decaying receptive fields. If the firing profile of border cells is exponential, and if the parameter controlling the spatial sensitivity of this profile differs across neurons, analogous to the differing relaxation times in Figure 2, then a population of border cells would code for the real Laplace transform of distance to the border. Applying the inverse transform to such a population would generate boundary vector cells (Lever, Burton, Jeewajee, O’Keefe, & Burgess, 2009), which have been observed in the subiculum and have been argued to drive classic hippocampal place fields (Burgess & O’Keefe, 1996). Analogously, trajectory coding cells and splitter cells observed in the EC and hippocampus can be understood as coding for the Laplace transform and inverse of functions over ordinal position—the sequence of movements leading up to the present (Frank, Brown, & Wilson, 2000; Wood, Dudchenko, Robitsek, & Eichenbaum, 2000). More broadly, this computational framework can be used to generate functions over a variety of spatiotemporal trajectories, which has been proposed to be a basic function the the MTL (Hasselmo, 2012; Dannenberg, Kelley, Hoyland, Monaghan, & Hasselmo, 2019). In this view, time cells and place cells are just two cases of a more general computational function (Kraus et al., 2013, 2015; Howard & Eichenbaum, 2015).

### Laplace transforms of time throughout the brain

If the brain contains a compressed record of the past (James, 1890; Husserl, 1966) as a neural representation across many different “kinds” of memory (Chater & Brown, 2008; Howard, Shankar, Aue, & Criss, 2015), then one might expect the existence of neurons with conjunctive receptive fields for what happened when across many different brain regions. Indeed, stimulus-specific time cells coding for what happened when have not only been found in regions believed to be important for episodic memory (MacDonald et al., 2013; Taxidis et al., 2018), but also regions that support working memory (Tiganj et al., 2018; Cruzado, Tiganj, Brincat, Miller, & Howard, 2019) and classical conditioning (Adler et al., 2012).

Because computational work using the Laplace transform has shown that a population of time cells can be constructed from temporal context cells (Shankar & Howard, 2012, 2013; Howard et al., 2014), we may speculate about the pervasiveness of this phenomenon across the brain. Perhaps temporal context cells may be found in other brain regions outside the EC to support this computation across separate regions of the brain. Alternatively, especially given the high sensory convergence within the EC, perhaps temporal context cells in the EC are utilized for generating time cells across the brain.

## Methods

### Subjects, training, and surgery

Two, male rhesus macaques (Macaca mulatta), 10 and 11 years old, and weighing 13.8 and 16.7 kg respectively, were trained to sit in a primate chair (Crist Instrument Company, Inc., Hagerstown, MD) and to release a touch-bar for fruit slurry reward delivered through a tube. The monkeys were trained to perform various tasks by releasing the touch-bar at appropriate times relative to visual stimuli presented on a screen. Magnetic resonance images of each monkey’s head were made both before and after surgery to plan and confirm implant placement. Separate surgeries were performed to implant a head post, then months later, a recording chamber, and finally a craniotomy within the chamber. All experiments were performed in accordance with protocols approved by the Emory University and University of Washington Institutional Animal Care and Use Committees.

### Electrophysiology

Each recording session, a laminar electrode array (AXIAL array with 13 channels, FHC, Inc.) mounted on a microdrive (FHC, Inc.) was slowly lowered into the brain through the craniotomy. Magnetic resonant images along with the neural signal were used to guide the penetration. Spikes and local field potentials were recorded using hardware and software from Blackrock, Inc., and neural data were sampled at 30 kHz. A 500 Hz high-pass filter was applied, as well as an electric line cancellation at 60 Hz. Spikes were sorted offline into distinct clusters using principal components analysis (Offline Sorter, Plexon, Inc.). Sorted clusters were then processed further by custom code in MATLAB to eliminate any data where minimum inter-spike interval was less than 0.001 s, and to identify any missed changes in signal (e.g., decreased amplitude in the waveform of interest, a new waveform appearing), using raster plots and plots of waveforms across the session for each neuron. When change in signal was identified, appropriate cuts were made to exclude compromised spike data from before or after a change point. 455 potential single neurons originally isolated in Offline Sorter were reduced to 357 single neurons. To further ensure recording location within the entorhinal cortex and identify from which cortical layers neurons were recorded, we examined each session’s data for the stereotypical, electrophysiological signature produced across entorhinal cortical layers at the onset of saccadic eye movement (Killian, Jutras, & Buffalo, 2012; Killian, Potter, & Buffalo, 2015; Meister & Buffalo, 2018). Recording sessions took place in both anterior and posterior regions of the entorhinal cortex. One recording session, which other electrode placement metrics suggest was conducted above the entorhinal cortex within the hippocampus, lacked this electrophysiological signature and was excluded from further analysis (8 single neurons were excluded from this session). No recording sessions showed the current source density electrophysiological signature of adjacent perirhinal cortex (Takeuchi, Hirabayashi, Tamura, & Miyashita, 2011) at stimulus onset.

### Experimental design and behavioral task

For all recordings, the monkey was seated in a dark room, head fixed and positioned so that the center of the screen (54.1 cm × 29.9 cm LCD screen, 120 Hz refresh rate, 1280 × 720 pixels, BenQ America Corp., Costa Mesa, CA) was aligned with his neutral gaze position and 60 cm away from the plane of the his eyes (equating to 25 screen pixels per degree of visual angle, or 1°/cm). Stimulus presentation was controlled by a PC running Cortex software (National Institute of Mental Health, Bethedsa, MD). Gaze location was monitored at 240 Hz with an infrared eye-tracking system (I-SCAN, Inc., Woburn, MA). Gaze location was calibrated before and during each recording session with calibration trials in which the monkey held a touch-sensitive bar while fixating a small (0.5°) gray square presented at various locations on the screen. The square turned yellow after a brief delay chosen uniformly from the interval from 0.40 s to 0.75 s. The monkey was required to release the bar in response to the color change for delivery of the fruit slurry reward. The subtlety of the color change forced the monkey to fixate the location of the small square to correctly perform those trials, therefore allowing calibration of gaze position to the displayed stimuli. Specifically, the gain and offset of the recorded gaze position were adjusted so that gaze position matched the position of the fixated stimulus. Throughout the session, intermittent calibration trials enabled continual monitoring of the quality of gaze position data and correction of any drift.

Before each image presentation, a crosshair (0.3°× 0.3°) appeared in one of eighteen possible screen locations. Once gaze position registered within a 3°× 3° window around the crosshair and was maintained within that spatial window for between 0.50 and 0.75 s (chosen uniformly), the image was presented. Images were large, complex images downloaded from the public photo-sharing website, Flickr (www.flickr.com). If necessary, images were resized by the experimenter for stimulus presentation (sized 30°× 15° for Monkey WR and 30°× 25° for Monkey MP). Monkeys freely viewed the image, and then the image vanished after gaze position had registered within the image frame for a cumulative 5 seconds. No food reward was given during image viewing trials. Each image presentation was followed by three calibration trials.

Image stimuli were unique to each session, and each image was presented twice within a session about 20 to 40 minutes apart. Images were presented in a block design so that novel and previously-seen images were presented throughout the session. Within a trial block, novel images (30 or 60) would first be shown, and then presented again in pseudorandom order. After completing a block of trials, a new block of trials would begin. In the first 16 sessions, a three-block design of 60 image presentations (30 novel) per block was used, with a total maximum of 180 image presentations per session. In the rest of the sessions (*n* = 41), there were a total maximum of 240 image presentations across two trial blocks (120 image presentations of which 60 were novel within each trial block).

### Analysis of Neural Firing Fields

In order to determine temporal firing fields, spikes were analyzed using a custom maximum likelihood estimation script run in MATLAB 2016a. We calculated model fits across all trials available for each particular neuron considering the time from 500 ms before image presentation to 5 s after image presentation. Fits of nested models were compared using a likelihood ratio test. In the present paper, we considered three models: a constant firing model, a model adding a Gaussian time term, and an ex-Gaussian model for which the time term was given by the convolution of a Gaussian and an exponential time term. The constant model, 

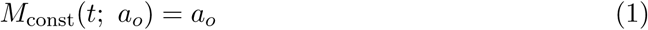

consisted of a single parameter *a*_*o*_ that predicted the constant probability of a spike at each time *t*.

The ex-Gaussian model describes the temporally-modulation of the firing field as the convolution of a Gaussian function with an exponentially decaying function: 

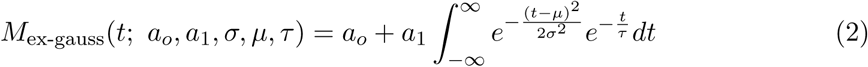

The ex-Gaussian distribution has been used extensively in studies of human response time data for many years (Ratcliff & Murdock, 1976). In the limit as *τ* → 0 the exponential function becomes a delta function and the result of the convolution in Eq. 2 is a Gaussian function. Similarly, in the limit as *σ* → 0 the Gaussian function becomes a delta function and the result of the convolution is an exponential function starting at *µ*. As such, this model is able to describe a range of peak firing times as well as varying degrees of skew (Fig. 1).

Two terms, *a*_*o*_ and *a*_1_ describe the contributions of the constant and time-modulated terms. Three parameters describe the shape of the temporally-modulated term (Figure S1). *µ* and *σ* describe the mean and standard deviation of the Gaussian distribution, which estimates the time that a neuron’s response maximally deviates from baseline, and the variability in that response time respectively. In the text, *µ* is referred to as the Response Time. *τ* measures the time constant of the exponential decay, and captures the time that a neuron has returned 63% of the way back to baseline. In the text, *τ* is referred to as the Relaxation Time.

To estimate parameters of Eq. 2 numerically we used an explicit form for the solution of the convolution in Eq. 2: 

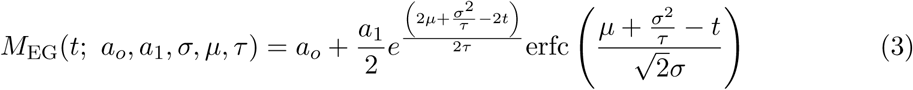

where erfc is the complementary error function. *µ* was allowed to take values between 0 and 5 s. *τ* was allowed to take values between 0 and 20 s. *σ* was allowed to take values between 0 and 1 s. Likelihood was estimated in a 5.5 s long window that included the 0.5 s prior to presentation of the image and the 5 s after presentation of the image.

We evaluated the models for each neuron *via* a likelihood ratio test and counted the number of neurons that were 1) better fit by the ex-Gaussian model at the 0.05 level, Bonferonni-corrected by the total number of 349 neurons, 2) changed their firing by at least 1 Hz, and 3) reached a firing rate of at least 3 Hz. In addition, we required that both even and odd trials for a neuron were significantly fit by the model, and that the those fits had a Pearson’s correlation coefficient greater than 0.4.

### Linear Discriminant Analysis (LDA)

An LDA classifier was used to decode time since onset of the image from the population using information from all neurons.

### LDA Implementation

Even and odd trials were used for training and testing respectively. The number of available trials varied for each neuron. To mitigate any problems from this, several steps were taken. First, four neurons with less than 30 trials each were entirely excluded from this analysis. Neurons with less than 200 trials were bootstrapped to 200 trials, while neurons with more than 200 trials were randomly down-sampled. Time was discretized into 0.25 s bins. For each bin of each trial, the firing rate was calculated across neurons. To avoid errors due to a singular covariance matrix, a small amount of uniform noise (between 0 and 1 × 10^−13^ Hz) was added to the firing rate in each time bin. The averaged firing rate of each time bin for each training trial across all neurons made up an element of the training data. The averaged firing rate of each time bin for each testing trial across all neurons made up an element of the testing data. LDA was implemented using the MATLAB function “classify.” This function takes in the training data, testing data, labels for the training data, and a selection of the method of estimation for the covariance matrix (the option “linear” was used) and returns a posterior distribution across bins for each test trial.

### Estimating the duration of temporal coding

To assess the quality of temporal information at different points within the interval, the LDA was repeated for successively fewer bins, at each step removing the earliest time bin. If time since presentation of the image can be decoded above chance using only information after time 𝒯, one can conclude that the population contained temporal information about time at least a time 𝒯 after presentation of the image. For each repetition the decoder was tested by training it on data with permuted time labels. We compared the absolute error of the actual data to the distribution generated from 1,000 permutations. The classifier’s performance was considered significantly better than chance if fewer than 10/1,000 permutations gave a better result than the unpermuted data (corresponding roughly to *p* < .01).

### Evaluating the distribution of temporal information over temporal context cells with different Relaxation Times

To assess the distribution of temporal information over temporal context cells as a function of their Relaxation Time, neurons with small Relaxation Times were progressively omitted from the LDA. First all temporal context cells were used (corresponding to a Relaxation Time Threshold of zero), then only cells with Relaxation Time longer than 0.125 seconds, then only cells with Relaxation Time longer than 0.25 seconds, and so forth. The longest Relaxation Time Threshold evaluated was 2.5 seconds. Performance was parametrized by averaging the absolute value of decoding error across all time bins. As a control, for each pseudo sub-population of cell, bins with permuted labels were also trained on and decoded from 1000 times.

### Stimulus sparsity analysis

To assess stimulus specificity in a way that facilitates comparison to previous human work, we followed the analysis method of (Mormann et al., 2008). In order to determine how many images a given neuron responded to, for each neuron we formed a distribution of baseline firing rates from the 500 ms prior to image onset on each trial. For each image, we took the trials in which they appeared and binned the firing rates of the 1 s following image onset into 19 overlapping bins, each of which were 100 ms long. We then compared these binned firing rates to the baseline distribution via a two-tailed Mann-Whitney U test, using the Simes procedure and a conservative significance threshold of *p* = 0.001. For each neuron, we counted the number of images that exceeded this threshold

### Population vector analysis of stimulus specificity

We constructed population vectors to evaluate the degree to which the entire population of entorhinal neurons was sensitive to the identities of visual images. For each repeated image, we created two population vectors, one corresponding to the first presentation and one corresponding to the second presentation. Each vector was created from the mean firing activity of all neurons recorded in a session during the 5 s of free viewing. Mean firing rates were normalized by each neuron’s maximum average firing rate so that firing rates ranged from 0 to 1. Only blocks where all images were presented twice were considered. In order to control for different block lengths between sessions, only the first 30 images presented in each block were used. All neurons that were recorded during first and second presentations of an image were included in this analysis (*N* = 332). The average number of simultaneously recorded neurons in a block was 8.51, with a standard deviation of 4.04, range 2-19. Similarity was measured by the cosine similarity of the two population vectors. We compared the cosine similarity of two presentations of the same image to the first presentation of one image and the second presentation of a different image. As a control we instead compared the population vector from the repetition of an image to the adjacent near-neighbors of the original image presentation. Near neighbors were required to be the first presentation of an image. Within session error bars represent the 95% confidence interval (Morey, 2008).

### Code and data availability

Code and data are available from the corresponding author upon request.

## Supplementary Information

### Time cells and temporal context cells: Two forms of temporal coding

The analyses in the main text differentiate two forms of temporal coding across populations of neurons. These two hypotheses are described in more detail here with simulated results for the primary empirical findings in the paper. Sequentially-activated time cells, like those observed in the hippocampus, are one possible form of temporal coding across a population of neurons. Exponentially-decaying temporal context cells are another possible form of temporal coding. The description of these ideal populations is based on prior theoretical work (Shankar & Howard, 2013).

The firing rate of the *i*th ideal temporal context cell a time *t* after the presentation of the image is given by 

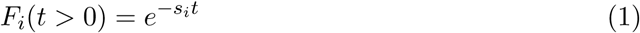

The rate constants *s*_*i*_ controls how fast ideal cell *i* decays. It can be shown that at time *t* after the image, the population codes the Laplace transform of the time in the past at which the image was presented (Shankar & Howard, 2012). In the simulations, the values of *s*_*i*_ were chosen such that *s*_*i*_/*s*_*i*+1_ was constant for all *i*, consistent with Weber-Fechner scaling. Equation 1 was used to generate the populations in Figure 1d and on the right of Figure S4.

The population of ideal sequentially activated time cells can each be described by values of *s* that correspond to those of the ideal temporal context cells. However, rather than rising immediately at time *t* = 0 and then decaying exponentially, the firing rate of these ideal time cells are given by 

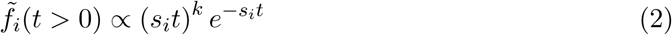

where *k* is a constant that was set to 4 in these figures. The properties of this population can be better understood by considering their activity as a function of 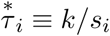, which has the units of time rather than rate. Then, 

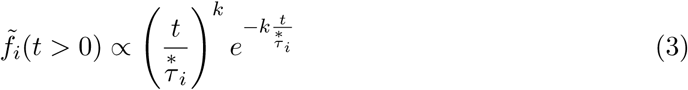

The time-dependence of the firing rate in Eq. 3 is the product of a growing polynomial term, 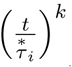 and a decaying exponential term, 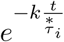 This product is zero at *t* = 0 and also zero as *t* → *∞* The product peaks somewhere in between that depends on the value of 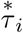 (it can be shown that this expression peaks precisely at 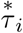) (Shankar & Howard, 2012). Because the time-dependence is a function of 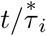, we can see that all such cells follow the same functional form of time-course. In the simulated results, the choice of *s*_*i*_ was the same as for the ideal temporal context cells.

### Relationship of µ to *τ* for populations of ideal time cells and ideal temporal context cells

Populations of cells generated by these two coding schemes, Eq. 1 and Eq. 3, will show different properties in their relationship between parameters *µ, σ* and *τ* estimated from the ex-Gaussian model. It is easy to convince oneself from Eq. 1 that cells generated from the exponentially-decaying term will have systematic changes in their value of *τ*_*i*_ (*τ*_*i*_ is just 1/*s*_*i*_) but there should not be a relationship between *τ*_*i*_ and *µ*_*i*_ or between *τ*_*i*_ and *σ*_*i*_. If the equations are precisely correct, the empirical values of *µ* and *σ* would result from the delay required for image information to reach the EC and initiate a response. That is, ideal temporal context cells would be expected to have about the same value of *µ* that is independent of *τ* and small values of *σ*. In contrast, for a population of ideal sequentially activated time cells because there are a range of peak times, we would expect to see a range of *µ*_*i*_. Moreover, because all of the cells have the same shape only rescaled with the peak time 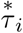 we would expect to see that *µ*_*i*_ correlates with both *τ*_*i*_ and *σ*_*i*_.

### Populations of time cells and of temporal context cells both contain information about time and have similar decoding properties

To illustrate the ability of both forms of temporal coding to carry information about time, we applied the LDA used on the data to two populations of simulated spiking data generated using the ideal equations shown above.

### Poisson spiking trains

To generate the simulated spiking data the following procedure was used. The ideal equation was treated as a normalized firing rate, were the maximum value (across all time constants) of the equation was normalized to 1. For each simulated millisecond, the probability of firing was set using this normalized simulated firing rate, a maximum firing rate selected for consistency with empirical spiking data, and a constant background firing rate. The probability of a spike within each millisecond is equal to the maximum firing rate multiplied by the normalized simulated firing rate plus a constant background firing rate. 1000 trials were simulated, 500 trials were used for training and 500 trials for testing.

### Simulated temporal context cells code for time with decreasing accuracy as time passes

Exponentially-decaying temporal context cells can be used to decode temporal information. The scale over which each neuron contributes maximally to decoding should be on the order of its time constant. Seventy simulated exponentially decaying cells were constructed using Eq. 1 with time constants ranging from 0.0125 s to 10 s, geometrically spaced with a ratio of about 1.1 between adjacent neurons. The simulated spiking data was then generated from this using a max firing rate of 20 Hz and a constant background firing rate of 20 Hz. This range of time constants means that the smallest time constant is less than 1/10 the duration of the bin size, and that the largest time constant is over twice the total duration being decoded. This reduces the possibility of any edge effects. Because there are fewer cells with slow time constants (because of the geometric spacing) decoding accuracy should go down with the passage of time. Figure S4 b and d shows results of the same LDA decoder used on the empirical data when applied to this simulated population of temporal context cells. As can be seen from the figure, temporal information decreased and error increased as a function of time. A linear regression of decoding error as a function of time showed a reliable slope, 0.13 ± 0.02 with *p* < 0.001, *R*^2^ = 0.73, as well as intercept 0.12 ± 0.05 with *p* < 0.05; *df* = 16. Thus the temporal information decreased as time elapsed.

### Simulated time cells code for time with decreasing accuracy as time passes

We also applied the LDA to a population of ideal time cells. These time cells were constructed using Eq. 2, with values of 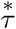 ranging from 0.05 s to 40 s, spaced geometrically with a ratio of approximately 1.1 (for a total of 70 simulated cells). In the case of time cells, their peak time is precisely equal to 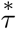, so their peaks range from 0.05 s to 40 s. The simulated spiking data was then generated from this using a max firing rate of 40 Hz and a constant background firing rate of 1 Hz. The width of the receptive fields expands with the peak time and because there are fewer neurons with peak times later in the delay, the decoding accuracy of this population of time cells should also go down with the passage of time. Figure S4a and c shows the results of the LDA applied to this set of simulated time cells. As can be seen from the figure, error increased as a function of time. A linear regression of decoding error as a function of time showed a reliable slope, 0.14 ± .03 with *p* < 0.001, *R*^2^ = 0.57, as well as intercept 0.22 ± 0.08 with *p* < 0.05; *df* = 16. Thus the temporal information decreases as time elapses.

Despite the fact that these two populations have different forms of temporal responsiveness, they both code information about time with similar properties. This is a natural consequence of the fact that the time cell population is just a linear transformation of the temporal context cell population (Shankar & Howard, 2012, 2013).

### Additional empirical results – EC neurons responded to many images

Previous work in humans has shown that, on average, EC neurons respond to a small number of stimuli, and that more selective neurons had longer latencies (Mormann et al., 2008). To assess the sparsity of monkey EC neurons and assess whether sparsity is confounded with Response Time, we determined the proportion of images for which a neuron’s firing rate changed after stimulus presentation (see “Stimulus Specificity” within the Methods for details). Supplementary Figure S7 shows the distribution of stimulus selectivity across neurons. On average, neurons showed a significant change in firing rate for 23% of presented stimuli. This corresponds to approximately 21 images, but note that the total number of images is not constant across neurons. Across temporal context cells (*n* = 109), we did not find a significant correlation between neural selectivity and Response Latency, *ρ* = −0.05, *p* = 0.6159. Although these results are not consistent with previous reports of stimulus specificity in monkey EC (Mormann et al., 2008), there are numerous methodological differences between the two experiments that may explain this discrepancy.

---

**Figure S1.**
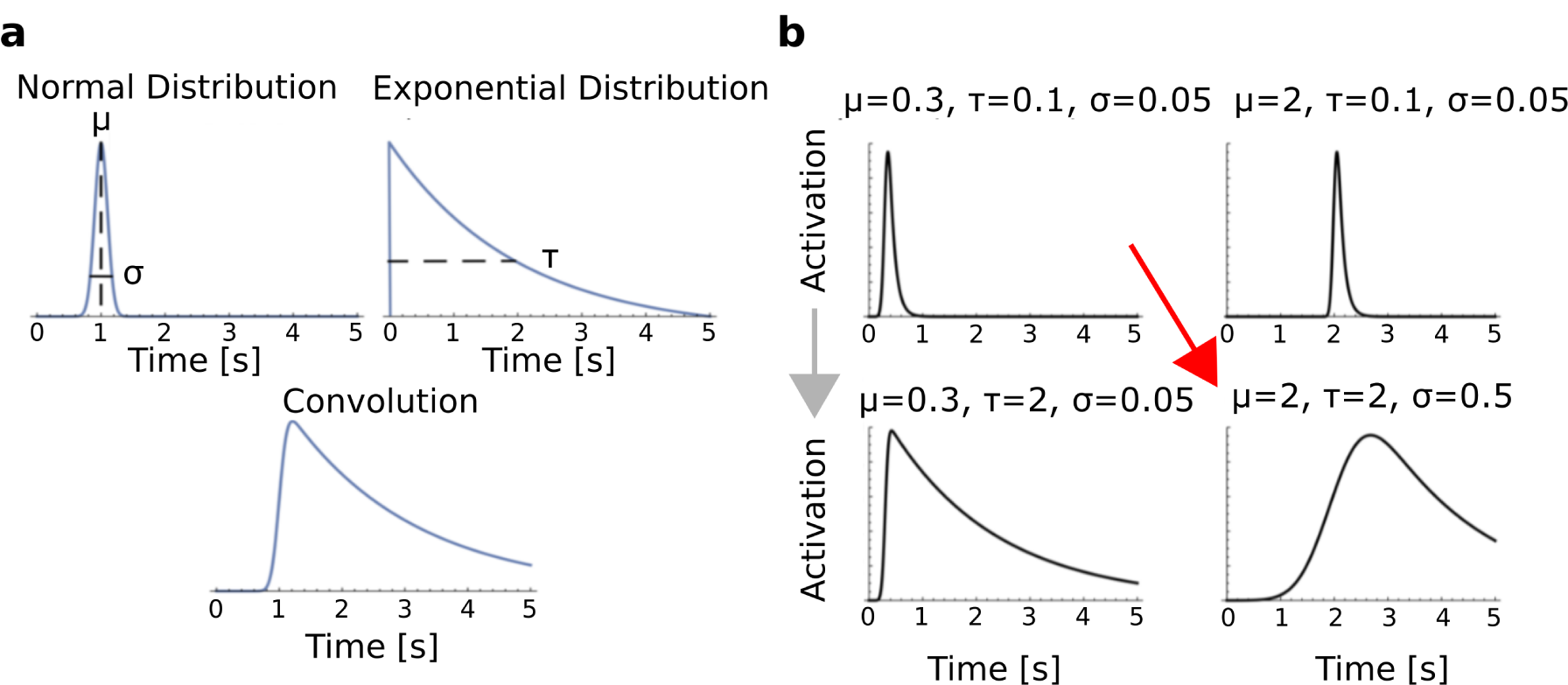
Modeling neuron responses using an ex-Gaussian distribution. **a**, The ex-Gaussian model of a neuron’s response (bottom) is formed by convolving a Gaussian distribution (with parameters *µ* and *σ*) with an exponential distribution (with parameter *τ*) (top). **b**, In the ex-Gaussian model, an increase in *µ* shifts the distribution to the right, an increase in *σ* widens the central peak of the distribution, and an increase in *τ* lengthens its decay rate. The gray and red arrows correspond to the parameter changes expected in the process of fitting the response of temporal context cells and sequentially activated time cells respectively.

**Figure S2.**
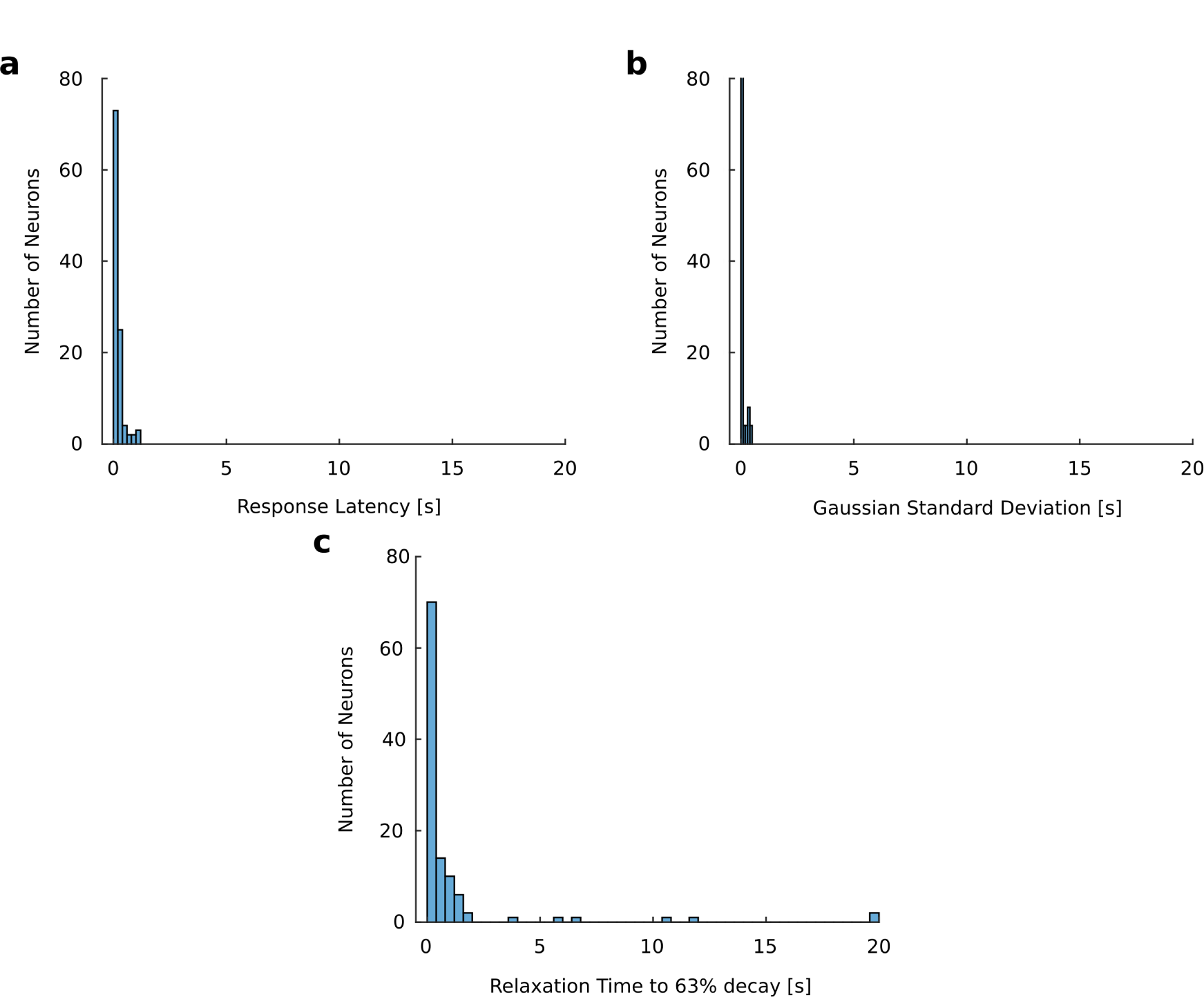
The distribution of individual parameters is consistent with temporal context cells. **a**, Neurons started responding (Response Latancy) shortly after image onset, and latencies did not cover the entire 5 s viewing period. **b**, Neurons did not show much deviation in their response latencies, with the majority having values of a few hundredths of a second. **c**, Most neurons decayed 63% of the way back to baseline firing (Relaxation Time) quickly, but some showed longer relaxation times that spanned the entire 5 s.

**Figure S3.**
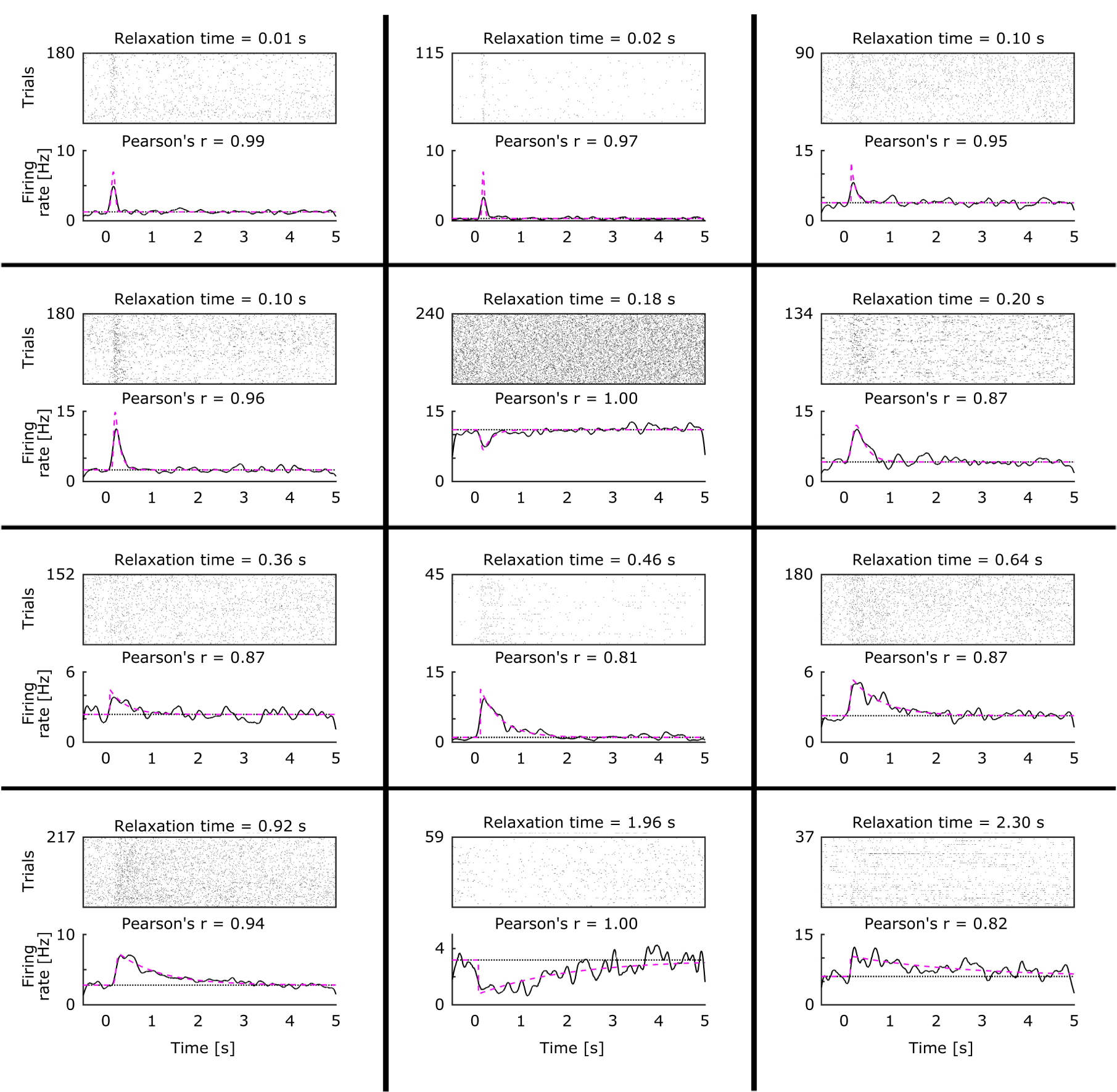
Additional examples of temporal context cells. Format is as in Figure 2a.

**Figure S4.**
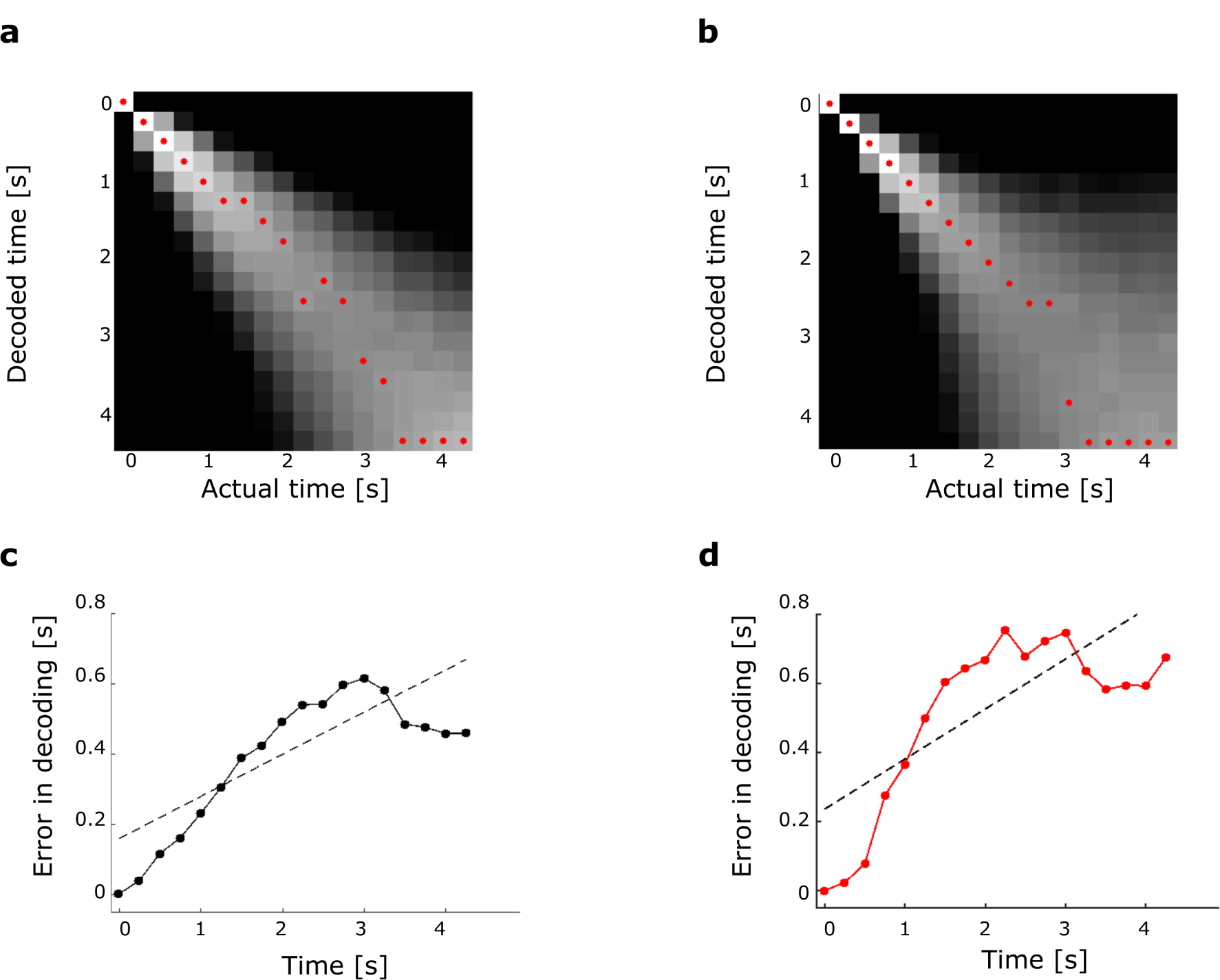
Time can be decoded from both ideal time cells and ideal exponentially decaying temporal context cells. Simulated noisy exponentially decaying temporal context cells (**a, c**) and simulated noisy time cells (**b, d**). Time is binned in 0.25 s bins. A linear decoder was trained on odd trials and tested on even trials. **a, b** The log of the average posterior probabilities of the classify function are averaged across trials for each time bin. The highest posterior decoded time for each actual time is marked with a red dot. Perfect decoding would manifest as a bright white diagonal with all of the red dots on the diagonal. The posterior distribution of the classifier shows increasing uncertainty in the decoded time for both populations. Colorbar and scaling are as in Figure 3a (maximum white correspond to probability 0.5 and greater, minimum black correspond to probability 0.02 and less, color is on log scale). **c, d** Averaged absolute value of decoding error. The decoding error goes up with time for both populations, as shown by the fitted regression line (dotted black line).

**Figure S5.**
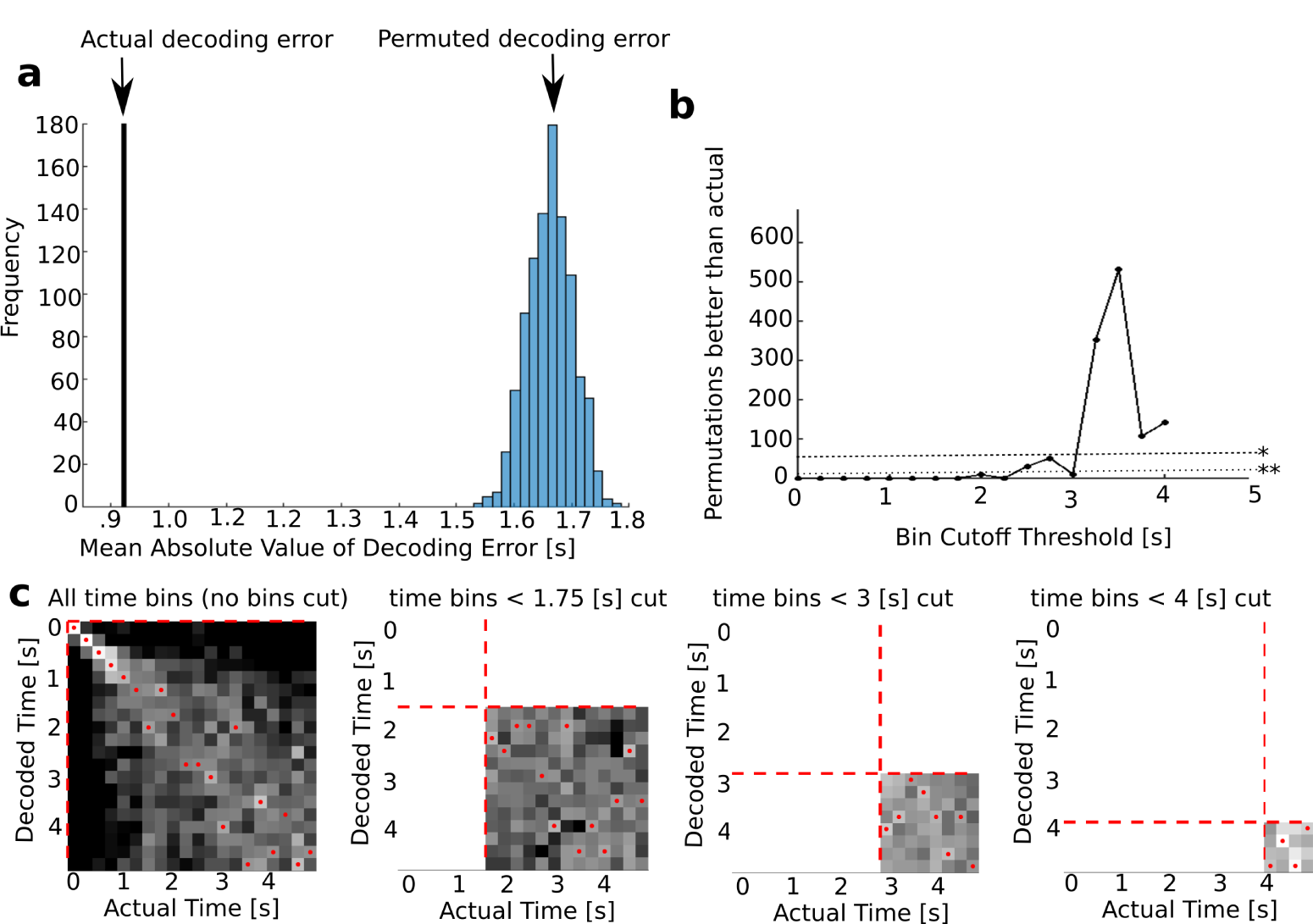
Time decoded from entorhinal firing showed a much lower decoding error than time decoded from permuted data. This is true even as bins are dropped, up to 2.25 seconds conservatively and 3 seconds less conservatively. **a**The histogram shows the distribution of mean absolute value of decoding error for the permuted data. In this analysis, the training data were the same as the data used in the actual decoder, except the true time labels were randomly permuted. The mean absolute value of decoding error for the original data (.93 s) is marked by a vertical line. This value is more than 18 standard deviations from the mean of decoding error of the permuted data. **b** Temporal information is present even in later time bins, the decoder performance is not merely an artifact of earlier time bins. This is demonstrated here by dropping the earlier bins and repeating the analysis against permuted data. As successive bins are dropped, permuted data begin to match the actual data in decoding accuracy. With 1000 permutations, the significance level is thus the number of permutations that matched or out performed the actual data divided by 1000 (thus the line marked by ‘**’ is significance at *p* < 0.01, the line marked by ‘*’ is significance at *p* < 0.05). Thus the decoder is always significantly better than chance at *p* < 0.01 up until 2.25 seconds (permutations remain below the line marked with ‘**’). The decoder is last significantly better than chance at *p* < 0.01 at 3 seconds. 500 bins is about the level of pure chance. Note that the accuracy varies some in that it appears that dropping noisy bins can relatively increase the decoder performance. Once the decoder is at chance it can vary substantially, but overall, dropping bins decreases performance. This suggests that temporal information is present out to at least 3 seconds. **c** Posterior distributions as bins are dropped. The leftmost sub-panel corresponds to no bins dropped, the next sub-panel corresponds to the first 1.75 seconds dropped the next sub-panel correspond to the first 2.5 seconds dropped, and the rightmost sub-panel corresponds to the first 3.75 seconds dropped) Colorbar and scaling are as in Figure 3a (maximum white correspond to probability 0.5 and greater, minimum black correspond to probability 0.02 and less, color is on log scale).

**Figure S6.**
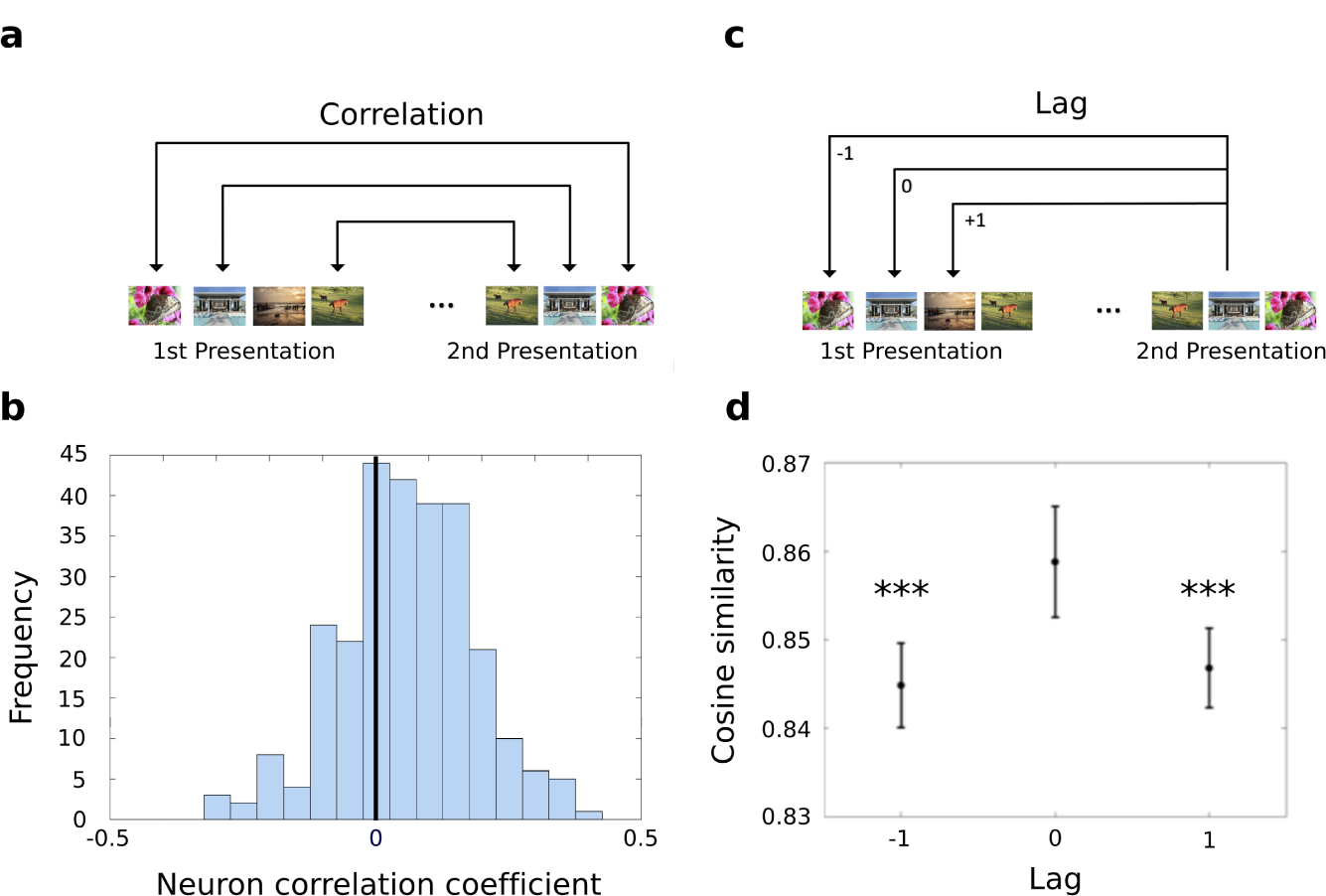
The entorhinal ensemble carries information about stimulus identity. **a**, We measured the firing rate of each unit to first and second presentations of each image. For each neuron that was recorded for an entire block of images (*n* = 270), we measured the correlation in firing rate across images using Kendall’s *τ*. **b**, The distribution of Kendall’s *τ* for all entorhinal neurons is shown. This distribution is reliably different from zero (*p* < 0.001, see text for details) indicating that entorhinal firing was sensitive to image identity. **c**, Schematic of a population similarity analysis. We measured the similarity of population response to the second presentation of each image to the first presentation of the same image (lag 0). As controls we also computed the similarity between the population response to the second presentation of an image and the responses to the images neighboring the first presentation of that image. Lag −1 refers to the similarity to the immediate predecessor of the image; lag +1 refers to the similarity to the immediate successor of the image. **d**, Cosine similarity of the population response to the second presentation of an image to the original presentation of an image (lag 0) and the images neighboring the original presentation of the image (lags −1 and +1) over 64 blocks of first and second presentations. The similarity to lag 0 is greater than the similarity to either of the neighboring images. *** indicates significance at the *p* < 0.001 level. Error bars correspond to the 95 % confidence interval of mean cosine similarity calculated across sessions.

**Figure S7.**
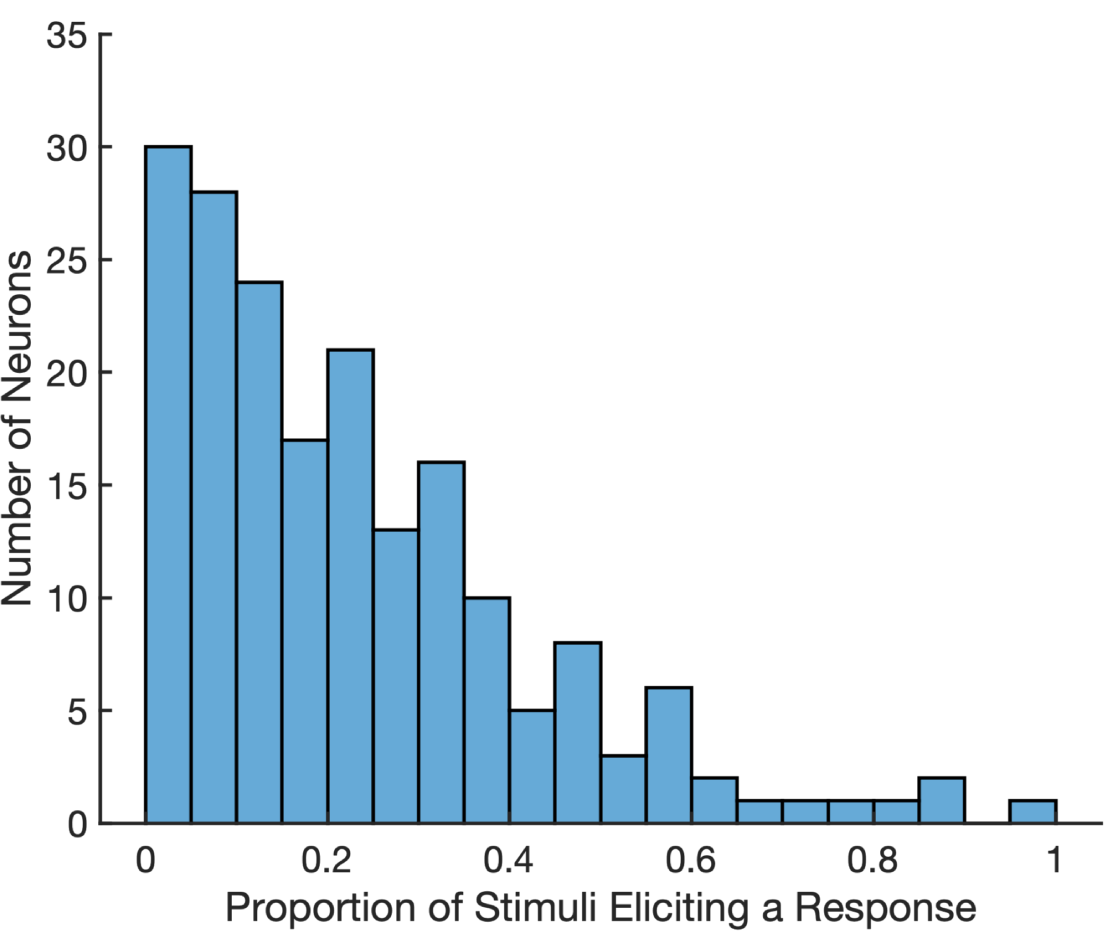
Entorhinal neurons showed different levels of stimulus specificity. A histogram of the proportion of images each neuron responded to. Image responsiveness was calculated by comparing a neuron’s firing rate during all presentations of an image to its baseline firing (see methods for details). Across the 349 neurons, most only responded to a few images, but a few neurons responded to the majority of images. On average, neurons responded to 23% of presented stimuli.

